# ATP13A4 upregulation drives the elevated polyamine transport system in the breast cancer cell line MCF7

**DOI:** 10.1101/2023.03.31.534207

**Authors:** Sarah van Veen, Antria Kourti, Elke Ausloos, Joris Van Asselberghs, Chris Van den Haute, Veerle Baekelandt, Jan Eggermont, Peter Vangheluwe

## Abstract

Polyamine homeostasis is disturbed in several human diseases, including cancer, which is hallmarked by increased intracellular polyamine levels and an upregulated polyamine transport system (PTS). So far, the polyamine transporters contributing to the elevated levels of polyamines in cancer cells have not yet been described, despite the fact that polyamine transport inhibitors are considered for cancer therapy. Here, we tested whether upregulation of candidate polyamine transporters of the P5B- transport ATPase family is responsible for the increased PTS in the well-studied breast cancer cell line MCF7 compared to the non-tumorigenic epithelial breast cell line MCF10A. We found that MCF7 cells present elevated expression of a previously uncharacterized P5B-ATPase ATP13A4, which is responsible for the elevated polyamine uptake activity. Furthermore, MCF7 cells are more sensitive to polyamine cytotoxicity, as demonstrated by cell viability, cell death and clonogenic assays. Importantly, overexpression of ATP13A4 WT in MCF10A cells induces a MCF7 polyamine phenotype, with significantly higher uptake of BODIPY-labelled polyamines and increased sensitivity to polyamine toxicity. In conclusion, we establish ATP13A4 as a new polyamine transporter in the human PTS and show that ATP13A4 may play a major role in the increased polyamine uptake of breast cancer cells. ATP13A4 therefore emerges as a candidate therapeutic target for anticancer drugs that block the PTS.

## 1. Introduction

Polyamines, such as spermidine and spermine, are ubiquitous organic polycations that are involved in a broad range of critical cellular functions, including gene expression, cell proliferation and differentiation. Intracellular polyamine levels are tightly regulated by the concerted action of biosynthesis, catabolism and polyamine transport. Whereas polyamine metabolism has been well characterized, the molecular players of mammalian polyamine transport system (PTS) are only just emerging. Our group recently characterized ATP13A2 and ATP13A3, two isoforms of the P5B-type transport ATPases (ATP13A2-5), as members of the PTS [1]. ATP13A2 works as a lysosomal polyamine exporter of endocytosed polyamines that contributes to cellular polyamine uptake, and is implicated in neurodegenerative disorders [2]. ATP13A3 has been genetically linked with pulmonary arterial hypertension, and is a major component of the mammalian PTS that is mutated and defective in the commonly used CHO-MG cell model, which is marked by a deficient PTS [3]. P5B ATPases share a high sequence similarity in the substrate binding site suggesting that also the ATP13A4 and ATP13A5 isoforms may function as polyamine transporters.

Polyamine homeostasis is disturbed in several human diseases, including neurodegeneration [2] and cancer [4]. Various cancers, including breast cancer, are hallmarked by increased intracellular polyamine levels as a consequence of upregulated polyamine biosynthesis and PTS activity [5-8]. Elevated polyamine content correlates with cancer aggressiveness and is linked to a poor prognosis for breast cancer patients [5]. This is also the case for high expression of ornithine decarboxylase (ODC, the key rate-limiting enzyme in polyamine synthesis) [6]. High levels of polyamines are associated with hyperproliferation and tumorigenesis through effects on various signaling cascades, such as the mitogen-activated protein kinase (MAPK) [4, 9, 10] and PI3K/AKT/mTOR pathways [11]. Estrogens, known to play a major role in breast cancer origin and progression, stimulate ODC expression and activity [12]. In addition, the breast cancer oncogenes MYC [13] and PI3KCA [14], increase the polyamine load by stimulating ODC-mediated polyamine synthesis and the PTS. So far, the contributing polyamine transporters remain unidentified, despite the fact that the PTS has become a promising target for cancer therapy. Polyamine transport inhibitors like AMXT-1501 that block cellular polyamine uptake [15] are currently being tested in clinical trials, among other in the context of breast cancer (clinicaltrials.gov, NCT05500508). In addition, polyamine analogues that utilize the PTS and disrupt polyamine metabolism are considered as a novel therapeutic strategy for breast cancer [12].

In the present study, we investigated the putative role of P5B ATPases in polyamine homeostasis in the human cell lines MCF7 and MCF10A, which are commonly used *in vitro* models in breast cancer research. MCF10A is a spontaneously immortalized, normal-like breast epithelial cell line derived from fibrocystic mammary tissue [16], whereas MCF7 is the most studied breast cancer cell line worldwide and was generated from pleural effusions from a patient with metastatic breast cancer [17]. MCF7 is estrogen receptor-positive and belongs to the luminal A breast cancer subtype [18]. Interestingly, we found higher expression of ATP13A4 in the MCF7 cells as compared to the MCF10A cells that contributes to the increased polyamine uptake and cytotoxicity, as well as activation of MAPK signaling. Our study is the first to characterize ATP13A4 as a polyamine transporter that plays a role in the PTS of breast cancer cells. Our work provides a stepping stone to further establish the potential of ATP13A4 as a therapeutic candidate in cancer.

## 2. Materials and Methods

### 2.1. Materials

The following reagents were purchased from Merck: dimethyl sulfoxide (DMSO; 276855), difluoromethylornithine (DFMO; D193), 4-methylumbelliferyl heptanoate (MUH; M2514), crystal violet (V5265), berenil (D7770), SigmaFast™ protease inhibitor (S8820), benzyl viologen dichloride (271845), putrescine dihydrochloride (P7505), spermine (S3256) and spermidine (S2626). Bovine serum albumin (BSA; 3854.3) was ordered from Carl Roth. N-(3-aminopropyl)cyclohexylamine (APCHA; sc-202715) and trans-4-methylcyclohexylamine (4MCHA; sc-272662) were obtained from Santa Cruz Biotechnology. TrypLE™ (12604021) and paraformaldehyde (J61899.AP) were ordered from ThermoFisher Scientific. The following antibodies were purchased from Cell Signaling Technology: phospho-JNK antibody (9251), phospho-AKT antibody (4058), phospho-ERK1/2 antibody (9101), HRP-linked anti-mouse IgG antibody (7076) and HRP-linked anti-rabbit IgG antibody (7074). Anti-GAPDH antibody (G8795) was purchased from Merck. Methylglyoxal bisguanylhydrazone (MGBG) and boron dipyrromethene (BODIPY)-conjugated polyamines [19] were supplied by Dr. P. van Veldhoven and Dr. S. Verhelst, respectively.

### 2.2. Preparation of polyamines and inhibitors

Polyamines were dissolved in 0.1 M MOPS (pH 7.0, KOH) to a final stock concentration of 500 mM (putrescine, spermidine) or 200 mM (spermine). The polyamine synthesis inhibitors 4MCHA and APCHA were diluted in DMSO to a final stock concentration of 500 mM. DFMO (500 mM), berenil (200 mM), benzyl viologen (200 mM) and MGBG (100 mM) were prepared in milliQ water to a final stock concentration as indicated between brackets. BODIPY-labeled polyamines were dissolved in 0.1 M MOPS-KOH (pH 7.0) to a final stock concentration of 5 mM.

### 2.3. Cell culture and lentiviral transduction

The cell lines MCF10A (CRL-10317) and MCF7 (HTB-22) were purchased from ATCC. MCF7 cells were cultured in Dulbecco’s Modified Eagle medium (DMEM; Gibco) supplemented with 10 % heat-inactivated fetal bovine serum (FBS; PAN BioTech), 1 % penicillin/streptomycin (Merck), 1 % non-essential amino acids (Merck) and GlutaMAX™ (Thermo Fisher Scientific). MCF10A cells were cultured in DMEM/F-12 medium (Thermo Fisher Scientific) supplemented with 5 % heat-inactivated horse serum (Merck), EGF (20 ng mL^−1^; Peprotec EC Ltd), hydrocortisone (0.5 μg mL^−1^; TCI Europe), cholera toxin (100 ng mL^−1^; Merck), insulin (10 μg mL^−1^; Merck), 1 % penicillin/streptomycin (Merck), 1 % non-essential amino acids (Merck) and GlutaMAX ™ (Thermo Fisher Scientific). Note that we used heat-inactivated FBS to deplete the polyamine oxidase activity, which may otherwise affect the supplemented fluorescent polyamine probes or polyamines. MCF7 and MCF10A cells were cultured at 37°C with 5% CO_2_. All cell lines were routinely tested for mycoplasma infection and found to be negative.

MCF10A cells with stable overexpression of human ATP13A4 WT or the catalytically dead mutant D486N were produced via lentiviral vector transduction. After lentiviral transduction, cells were selected with 1 μg mL^−1^ puromycin.

### 2.4. RT-qPCR

mRNA expression levels of ATP13A2-4 were quantified by RT-qPCR analysis. Therefore, we extracted RNA from 1.0 × 10^6^ cells using NucleoSpin RNA plus Kit (740984, Macherey-Nagel) according to the manufacturer’s instructions. The concentration and purity of the RNA samples were measured using a Nanodrop spectrophotometer (Thermo Fisher), followed by conversion of RNA into cDNA using the RevertAid H Minus First Strand cDNA Synthesis Kit (K1631, Thermo Fisher). The cDNA was then subjected to SYBR Green-based qPCR with gene-specific primer pairs (Table 1). β-actin was used as a reference gene. The analysis was performed using Light Cycler machine (Roche) and the cycling conditions were as follows: 10 min at 95°C, 50 cycles: 10 s at 95°C, 30 s at 55°C, 1 min at 95°C and 1 min at 55°C. Melting curves were analyzed from 55 to 95°C. Finally, the mean Cq values were determined.

**Table 1.**
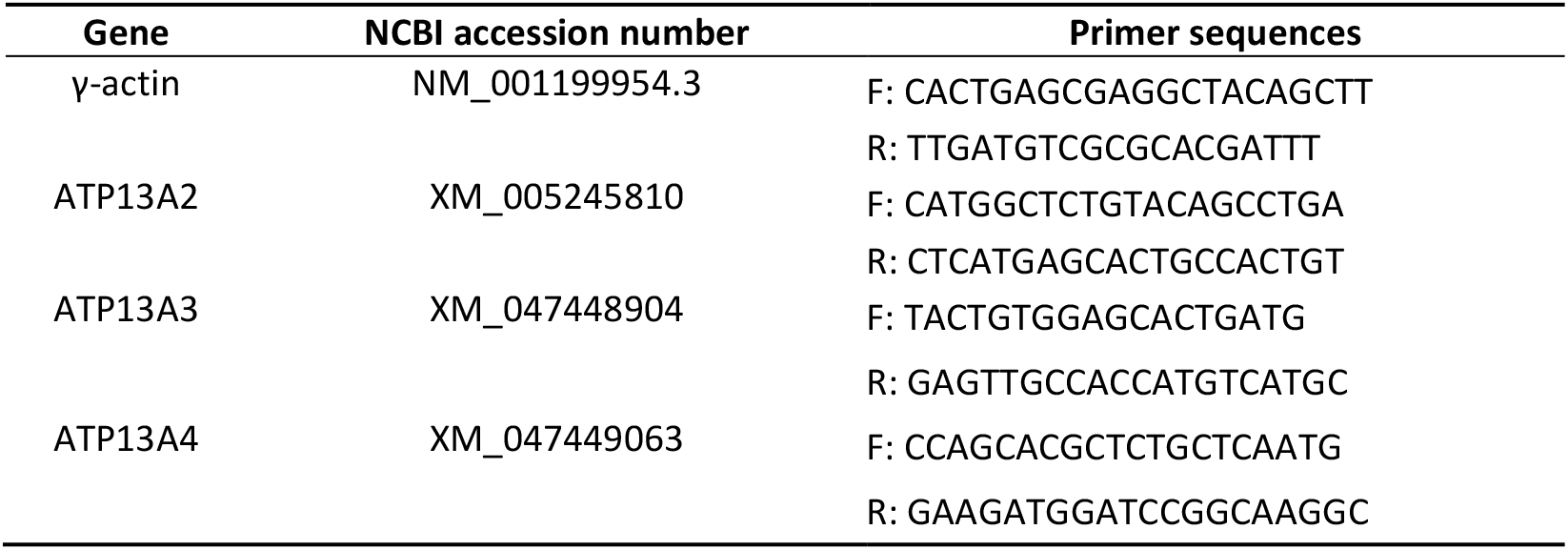
Gene accession numbers and primer sequences.

### 2.5. BODIPY-polyamine uptake

For flow cytometry-based measurement of BODIPY-polyamine uptake, cells were seeded in 12- well plates at a density of 1.0 × 10^5^ cells per well. The next day, cells were incubated for 2 h with 1 μM BODIPY-conjugated polyamines, *i.e*. putrescine, spermidine and spermine. To evaluate the effect of inhibitors of polyamine uptake or synthesis, the cells were pre-treated with 1 mM benzyl viologen (90 min) or 10 μM MGBG (30 min), respectively. Afterwards, cells were washed and resuspended in PBS supplemented with 1 % BSA. Uptake was measured by recording the mean fluorescence intensities (MFI) of 10,000 events using a flow cytometer (ID7000 spectral cell analyzer, Sony).

For confocal microscopy-based analysis of BODIPY-polyamine uptake, cells were seeded on cover slips in 12-well plates at a density of 5.0 × 10^4^ cells per well. The next day, cells were incubated for 2 h with 1 μM BODIPY-conjugated polyamines, washed in PBS and fixed in 4% paraformaldehyde for 30 min at 37°C. Thereafter, cells were washed in PBS and stored at 4°C. The next day, cells were stained with DAPI to visualize the nucleus. Cover slips were mounted on slides and images were acquired using an LSM880 confocal microscope (Zeiss) with a 63x objective.

### 2.6. Measurement of cell viability and cell death

Cells were seeded in 96-well plates (1.0 × 10^4^ cells per well) and the following day, cells were treated with increasing concentrations of the different polyamines and inhibitors for 24 h to 1 week, as indicated. For the 1-week treatments, we plated 5.0 × 10^3^ cells per well, and every 48 h we replaced the medium containing the respective inhibitor. Following the incubation period, we assessed cell death (ToxiLight assay, Lonza, LT07-117) and cell viability (MUH cell viability assay [2]).

### 2.7. Colony formation assay

Cells were seeded in a 12-well plate (MCF7: 5.0 × 10^3^ cells per well, MCF10A: 1.0 × 10^3^ cells per well) and were treated the next day with indicated concentrations of spermidine or spermine. Every 5 days the media was discarded and new media containing the treatment of interest was added. After 14 days, the media was removed and cells were washed with PBS. The cells were fixed with methanol for 15 min and then stained with 0.01 % (w/v) crystal violet solution in milliQ water for 1 h. Afterwards, the plates were carefully rinsed with distilled water and left to dry at room temperature. An ordinary scanner was used to take images from the 12-well plates. To quantify the staining (relative to the untreated control condition), crystal violet was solubilized from the stained colonies using 10 % acetic acid (1 h incubation with gentle rocking, room temperature) and absorbance was read at 590 nm, as described in [20].

### 2.8. Western blotting

Cells were seeded in 10 cm plates (1.0 × 10^6^ cells per plate) and the following day, cells were treated with spermidine (100 μM) or spermine (100 μM) for indicated times. Thereafter, cells were dissociated from the plates using TrypLE™ and lysed in RIPA buffer supplemented with protease and phosphatase inhibitors (A32957, Thermo Fisher Scientific). Proteins in cell lysates (20-40 μg) were separated on precast NuPAGE™ 4-12% Bis-Tris gels (Invitrogen) using MES (Figure 4, S5) or MOPS (Figure 3A) running buffer (Life Sciences), followed by transfer onto polyvinylidene fluoride (PVDF) membranes (Millipore) according to the manufacturer’s instructions. After blocking in TBS-T (50 mM Tris-HCl, 150 mM NaCl, pH 7.5, 0.1 % Tween-20 (Sigma)) supplemented with 5 % non-fat dry milk, blots were incubated with primary antibodies (1/1,000 dilution in TBS-T with 1 % BSA) and horseradish peroxidase (HRP)-conjugated anti-rabbit secondary antibodies (1-2 h; 1/1,000 dilution in TBS-T with 1 % BSA). We used primary antibodies directed against phospho-JNK, phospho-ERK1/2, phospho-AKT, ATP13A4 and GAPDH. The rabbit polyclonal anti-ATP13A4 antibody was home-made and raised against the epitope ^1182^VSYSNPVFESNEEQL. Blots were incubated with phospho-specific antibodies overnight at 4°C, whereas the incubation time for anti-ATP13A4 and anti-GAPDH antibodies was 1 h at room temperature. Protein expression was detected using enhanced chemiluminescence (ECL) substrate (Pierce) and the Bio-Rad ChemiDoc MP imaging system. Quantification was performed with ImageJ software (https://imagej.net/ij/index.html).

### 2.9. Statistical analysis

Data analysis was performed using GraphPad Prism 9 (La Jolla, CA, USA). Figure legends cover the type of statistical test used. Data are presented as the mean ± s.e.m. and in the bar graphs, individual points are also shown. Each experiment was repeated at least three times, except for Figure S5 (N = 2).

## 3. Results

The tumorigenic human breast cancer cell line MCF7 presents upregulated PTS activity compared to the non-tumorigenic epithelial breast cell line MCF10A [21], but the molecular players responsible for this phenotype remain unknown. Here, we hypothesized that isoforms of the P5B transport ATPases may be implicated.

### 3.1. MCF7 cells show enhanced PTS activity compared to MCF10A cells

First, we confirmed via cellular uptake experiments with the fluorescent BODIPY-labeled putrescine, spermidine and spermine, that MCF7 cells present an elevated PTS compared to MCF10A cells (Figure 1A-D). Flow-cytometry-based uptake experiments revealed an over three-fold higher uptake of BODIPY-labeled putrescine (Figure 1A) and spermidine (Figure 1B) in MCF7 cells compared to MCF10A cells, whereas the uptake of BODIPY-spermine was over ten-fold higher (Figure 1C). These findings were confirmed by confocal fluorescent microscopy (Figure 1D). Whereas we observed strong fluorescence in the MCF7 cells incubated with BODIPY-polyamines, only weak fluorescent intensity was detected in the MCF10A cells. The difference between the cell lines was highest for BODIPY-labelled spermine (Figure 1D, bottom panels). Furthermore, to prove that the fluorescent probes enter the cell via the PTS, we investigated the impact of methylglyoxal bis-(guanylhydrazone) (MGBG), a spermidine analogue that is taken up via the PTS competing with the cellular uptake of polyamines, and benzyl viologen (BV), a commonly used PTS inhibitor [3, 21, 22]. Both MGBG (10 μM, 30 min pre-treatment) and BV (1 mM, 90 min pre-treatment) significantly reduced the uptake of BODIPY-labeled putrescine (Figure 1A, Figure S2A), spermidine (Figure 1B, Figure S2B) and spermine (Figure 1C, Figure S2C) in the MCF7 cells, but not in MCF10A cells. These findings suggest that, in sharp contrast to the MCF7 cells, MCF10A cells only have minimal PTS activity.

**Figure 1.**
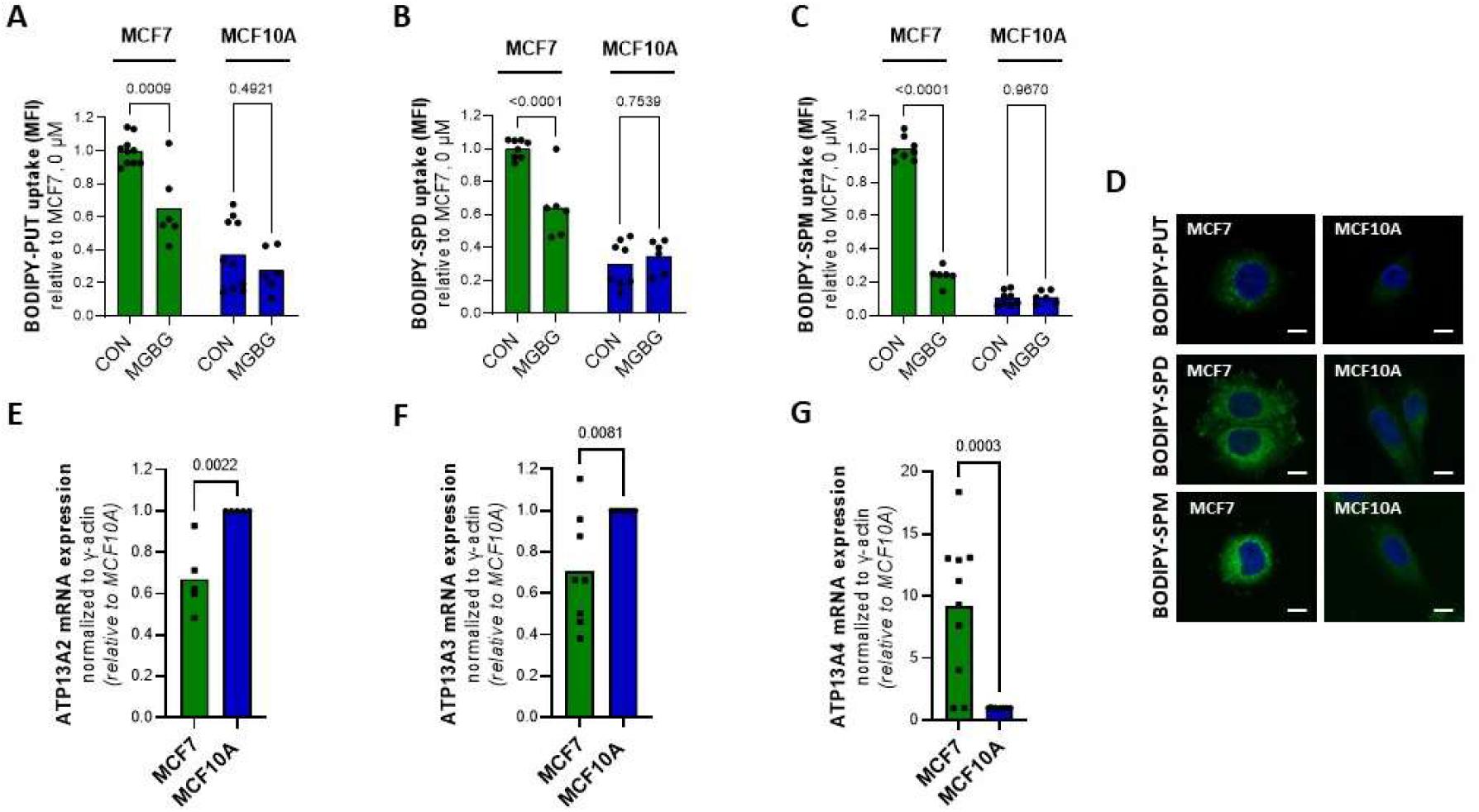
MCF7 cells exhibit an upregulated PTS and higher expression of ATP13A4 compared to MCF10A cells. Cellular uptake of BODIPY-labeled putrescine (BODIPY-PUT) (**A, D**), spermidine (BODIPY-SPD) (**B, D**) and spermine (BODIPY-SPM) (**C, D**) (1 μM, 2 h) in MCF7 *versus* MCF10A cells was assessed via flow cytometry (**A-C**) or confocal microscopy (**D**). Representative confocal microscopy images are shown of three independent experiments. Scale bar, 10 μm. MFI, mean fluorescence intensity. Expression of ATP13A2 (**E**), ATP13A3 (**F**) and ATP13A4 (**G**) was evaluated at the mRNA level via qPCR. mRNA expression was normalized to β-actin. Data in the bar graphs are presented as the mean of minimal three independent experiments with individual data points shown. Statistical significance was determined by one-way ANOVA with Šídák’s multiple comparisons test (**A-C**) or unpaired two- tailed t-test (**E-G**). *p* values are depicted in the graphs.

To further assess the importance of the PTS, we compared the sensitivity of the MCF7 and MCF10A cell lines to BV by measuring cell viability using the MUH assay. Interestingly, the MCF7 cells demonstrated increased cytotoxicity following 48 h treatment with BV, compared to MCF10A cells (Figure S2D), indicating that the MCF7 cells rely more on a functional PTS to maintain their overall health.

Altogether, our data show that the PTS is upregulated in MCF7 cells *versus* MCF10A cells.

### 3.2. The P5B ATPase ATP13A4 is upregulated in MCF7 versus MCF10A cells

To test whether specific P5B ATPase isoforms may play a role in MCF7 cells, we evaluated the relative expression of ATP13A2-5 between both cell lines by mining the ARCHS4 web resource of RNA-seq data [23] as a starting point (Figure S1). All P5B ATPases are expressed in MCF7 and MCF10A cells (Figure S1B) with ATP13A2 and ATP13A3 presenting similar high expression levels in both cell lines, whereas ATP13A4 and ATP13A5 are expressed to a lesser extent, but with higher expression in MCF7 *versus* MCF10A cells. Of note, MCF7 appears amongst the highest ATP13A4 expressing cell lines (Figure S1A).

To corroborate this finding, we performed qPCR to assess the mRNA levels of the P5B ATPases ATP13A2-5 in MCF7 and MCF10A cells (Figure 1E-G) and confirmed that MCF7 cells present significantly higher ATP13A4 mRNA levels than MCF10A cells, with an almost tenfold difference in expression (Figure 1G). Surprisingly, the expression of ATP13A2 and ATP13A3 was significantly lower in MCF7 than in MCF10A cells (Figure 1E-F), whereas we were unable to detect ATP13A5 mRNA in both cell lines (data not shown).

Based on this comparison, ATP13A4 emerges as a candidate polyamine transporter that may be responsible for the increased PTS in MCF7 cells.

### 3.3. MCF7 cells are more sensitive to polyamine cytotoxicity

At high concentrations, polyamines become toxic and here, we investigated the relative cytotoxicity of exogenous polyamines in the MCF7 and MCF10A cell lines as a complementary readout of the PTS activity (Figure 2).

**Figure 2.**
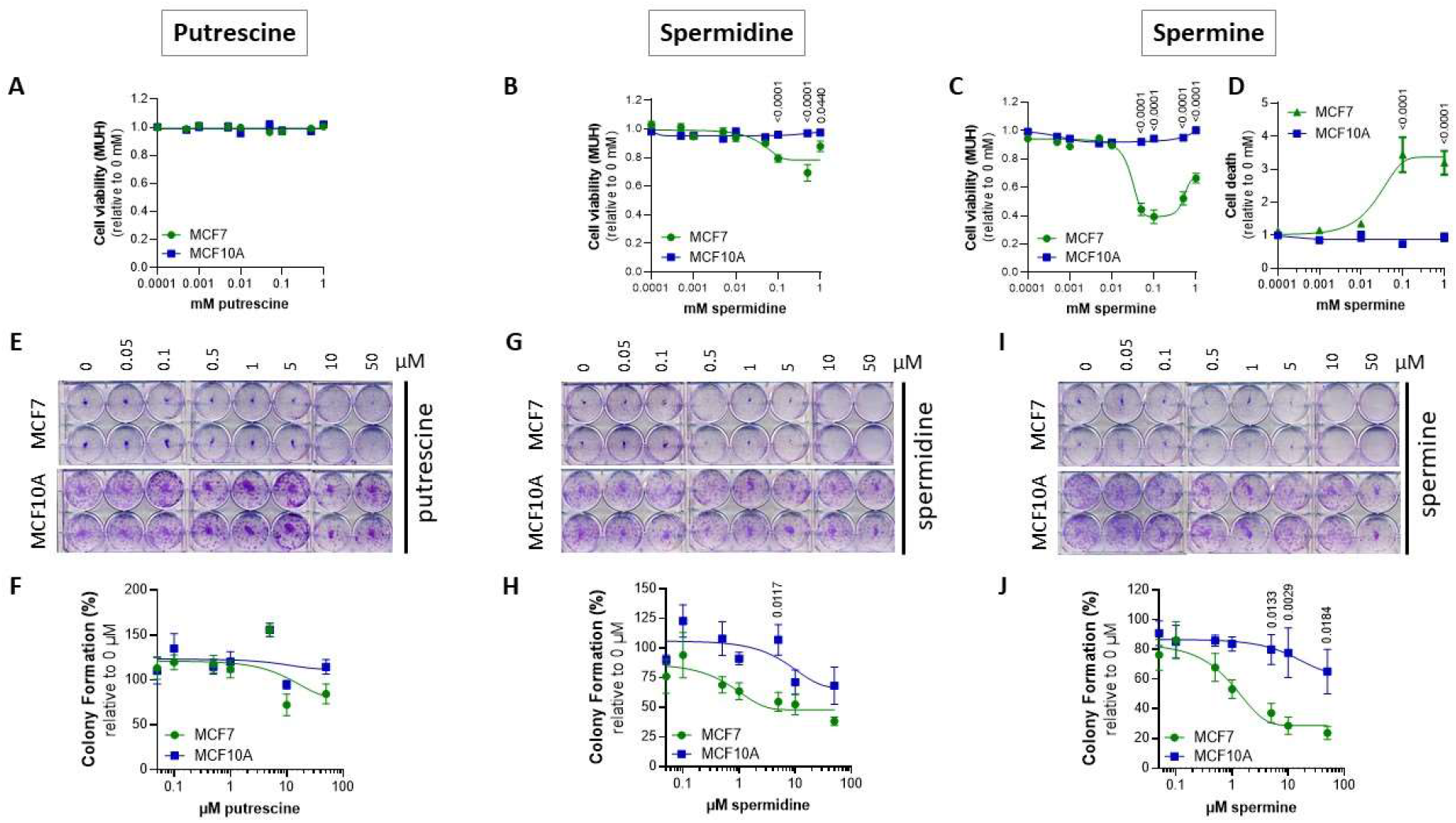
MCF7 cells are more sensitive to polyamine cytotoxicity than MCF10A cells. MCF7 and MCF10A cells were treated for 24 h (**A-D**) or 2 weeks (**E-J**) with indicated concentrations of putrescine (PUT) (**A**), spermidine (SPD) (**B**) and spermine (SPM) (**C-D**). Cell viability and cell death were assessed via MUH assay (**A-C**) and Toxilight (**D**) assays, respectively. Colony formation ability was evaluated via clonogenic assays (**E, G, I**) and quantified based on crystal violet absorbance (590 nm) (**F, H, J**). **E, G** and **I** depict representative pictures of minimal three independent experiments. Data were normalized to untreated control for each cell line, and are presented as the mean ± s.e.m. of minimal three independent experiments. Statistical significance was determined by two- way ANOVA with Šídák’s multiple comparisons test. *p* values are depicted in the graphs, non-significant values are not indicated.

First, we treated MCF10A and MCF7 cells for 24 h with increasing concentrations of putrescine, spermidine and spermine to assess their impact on cell viability. As shown in Figure 2A-C, MCF10A cells were insensitive to the tested polyamines in line with a low PTS activity. In contrast, MCF7 cells showed lower viability following treatment with spermine (Figure 2C) and to a lesser extent spermidine (Figure 2B), but not putrescine (Figure 2A). In addition, we assessed cell death following the exposure to spermine with the ToxiLight cytolysis assay that measures adenylate kinase, an enzyme that is released by dying cells. High levels of spermine led to increased cell death in MCF7 cells, whereas no cell death was observed in MCF10A cells (Figure 2D). These data are in line with the cell viability results (Figure 2C) implying that the reduced cell viability in MCF7 cells following spermine treatment is due to an increased cell death.

Furthermore, we cross-validated the polyamine cytotoxicity data via colony formation assays (Figure 2E-J) evaluating the impact of each polyamine on the ability of a single cell to proliferate and grow into a colony. The polyamines spermidine (Figure 2G-H) and especially spermine (Figure 2I-J) inhibit colony formation more severely in MCF7 cells than in MCF10A cells, which appear more resistant. Putrescine did not differentially impact clonogenic ability (Figure 2E-F).

To examine the role of polyamine metabolism, we compared the effect of pharmacological inhibitors of the polyamine metabolic pathway (Figure S3A) on the cell viability of MCF7 and MCF10A cells, which have previously been considered for cancer therapy. MCF7 cells appeared more sensitive to inhibition of the polyamine synthesis pathway by treating the cells for one week with DFMO (ornithine decarboxylase inhibitor), 4MCHA (spermidine synthase inhibitor) or APCHA (spermine synthase inhibitor), and less sensitive to MGBG (S-adenosyl-L-methionine-decarboxylase inhibitor); although the differences are mild (Figure S3B-E). In addition, after one week incubation, MCF7 cells are also slightly more sensitive to berenil, which inhibits SAT1, a key enzyme in polyamine catabolism (Figure S3F). The overall comparable sensitivity to polyamine metabolism inhibitors indicates that polyamine metabolism may not be very different between both cell lines.

All in all, our findings demonstrate that MCF7 cells are more sensitive to polyamine cytotoxicity, in particular spermine, than MCF10A cells, whereas polyamine metabolism blockers do not have a strong differential impact.

### 3.4. Overexpression of ATP13A4 in MCF10A cells induces an MCF7 polyamine phenotype

To investigate the causal role of ATP13A4 in the differential polyamine phenotypes of MCF7 and MCF10A cells, we overexpressed ATP13A4 in MCF10A to recapitulate the polyamine phenotype of MCF7 cells. Therefore, we generated stable MCF10A cells with overexpression of ATP13A4 via lentiviral transduction (WT or a transport dead mutant D486N that is mutated in the catalytic site for autophosphorylation) (Figure 3A).

**Figure 3.**
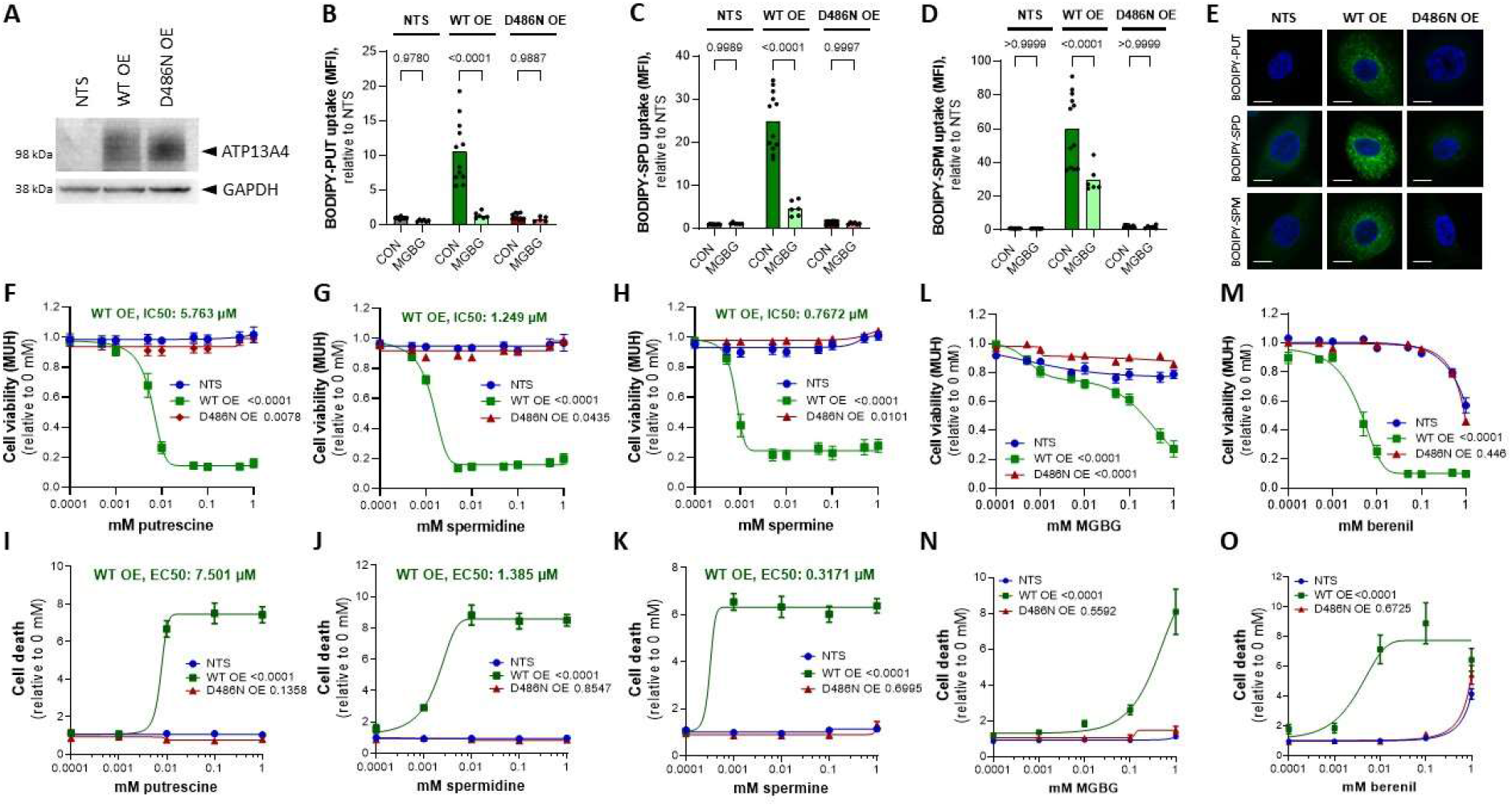
Overexpression of ATP13A4 WT in MCF10A cells induces an MCF7 polyamine phenotype. **A**. Immunoblot depicting ATP13A4 overexpression levels in MCF10A cells (non-transduced, NTS; ATP13A4 WT overexpression, WT OE; ATP13A4 D486N mutant overexpression, D486N OE). Cellular uptake of BODIPY-labeled putrescine (BODIPY-PUT) (**B, E**), spermidine (BODIPY-SPD) (**C, E**) and spermine (BODIPY-SPM) (**D-E**) (1 μM, 2 h) was assessed via flow cytometry (**B-D**) or confocal microscopy (**E**). Representative confocal microscopy images are shown of three independent experiments. Scale bar, 10 μm. MFI, mean fluorescence intensity. Cells were treated for 24 h with indicated concentrations of putrescine (**F, I**), spermidine (**G, J**), spermine (**H, K**), MGBG (**L, N**) or berenil (**M, O**) prior to cell viability (MUH; **F, G, H, L, M**) or cell death (Toxilight; **I, J, K, N, O**) assays. Data were normalized to untreated control for each cell line and are presented as the mean of minimal three independent experiments, with individual data points shown (**B-D**) or ± s.e.m. (**F-O**). Statistical significance was determined by one-way ANOVA with Šídák’s multiple comparisons test (**B-D**) or by two-way ANOVA with Dunnett’s multiple comparisons test (F-O, compared to NTS). p values are depicted in the graphs.

Interestingly, the uptake of all BODIPY-labeled polyamines was significantly increased in MCF10A cells as a consequence of the ATP13A4 WT overexpression; with the largest uptake effect on BODIPY- spermine (60-fold) over BODIPY-spermidine (25-fold) and BODIPY-putrescine (10-fold) as compared to non-transduced MCF10A cells (Figure 3A-D). In addition, the higher cellular uptake of BODIPY- polyamines was not observed with the D486N mutant, demonstrating that the catalytic transport activity of ATP13A4 is required (Figure 3B-D). The higher uptake of BODIPY-labeled polyamines in MCF10A cells that overexpress ATP13A4 WT was confirmed by confocal microscopy (Figure 3E). Furthermore, MGBG significantly reduced the uptake of BODIPY-labeled putrescine (Figure 3B), spermidine (Figure 3C) and spermine (Figure 3D) in MCF10A cells with overexpression of ATP13A4 WT, but not in non-transduced or D486N expressing MCF10A cells. Altogether, we present here the first evidence that ATP13A4 functions as a polyamine transporter within the mammalian PTS that may present a broad polyamine specificity.

To complement our findings with polyamine uptake in the MCF10A cell lines, we examined polyamine cytotoxicity, which in MCF7 cells is correlated with the elevated PTS (Figure 2). ATP13A4 WT overexpression strongly sensitized MCF10A cells to putrescine (Figure 3F, I), spermidine (Figure 3G, J) and spermine (Figure 3H, K), as evidenced by reduced cell viability and increased cell death. Of note, spermine exhibited the lowest IC50 and EC50 values, followed by spermidine and putrescine. These phenotypes are not observed in non-transduced MCF10A cells or cells overexpressing the catalytically inactive D486N mutant (Figure 3F-K), which confirms that the polyamine transport activity plays a role. In addition, overexpression of ATP13A4 WT, but not the D486N mutant, sensitized MCF10A cells to MGBG toxicity in line with the increased PTS activity (Figure 3L, N). Remarkably, these cells also become highly sensitive to SAT1 inhibition via berenil, pointing to a protective role of SAT1- mediated polyamine degradation in these cells (Figure 3M, O). Of note, the cytotoxicity phenotypes are much more pronounced in the MCF10A cells than in the MCF7 cells, most likely as a consequence of the high ATP13A4 overexpression levels. This may also explain the observed putrescine toxicity, which was not observed in the MCF7 cells. Lastly, we did not observe a reducing effect of BV on BODIPY-polyamine uptake in ATP13A4 overexpressing MCF10A cells or an impact on viability, indicating that BV may not be a direct inhibitor of ATP13A4 (Figure S4).

Together, our findings with polyamine uptake and cytotoxicity assays indicate that overexpressing ATP13A4 WT in MCF10A cells confers a polyamine uptake phenotype resembling MCF7 cells with high endogenous ATP13A4 expression, indicating that ATP13A4 is the driver for the increased PTS in MCF7 breast cancer cells.

### 3.5. ATP13A4 overexpression triggers activation of the JNK signaling pathway in MCF10A cells

One of the most commonly altered pathways driving breast cancer cell progression is the PI3K/AKT/mTOR signaling cascade [24], whereas also the N-terminal c-Jun kinase (JNK) [24] and extracellular signal-regulated kinase (ERK) [24] in the MAPK pathway have been implicated [10]. Moreover, interplay between these signaling pathways and polyamines has been described before [4, 14, 25]. Therefore, we investigated whether these signaling pathways are altered between MCF7 and MCF10A cells and respond to toxic polyamine supplementation. At a basal level, phosphorylated JNK and ERK1/2 were hardly detectable in MCF10A cells compared to MCF7 cells, while phosphorylated AKT levels were much higher in MCF10A cells (Figure S5). The polyamines spermidine and spermine did not cause significant changes in phosphorylation of JNK, ERK or AKT over time (Figure S5A-H), although modest effects on JNK and AKT phosphorylation were observed (Figure S5B, D, F, H). However, MCF10A cells overexpressing ATP13A4 WT (but not D486N) and treated with spermidine (Figure 4A-B) or spermine (Figure 4C-D) presented a fast and robust upregulation of JNK phosphorylation, which is reminiscent of the basal JNK phosphorylation phenotype of MCF7 cells.

**Figure 4.**
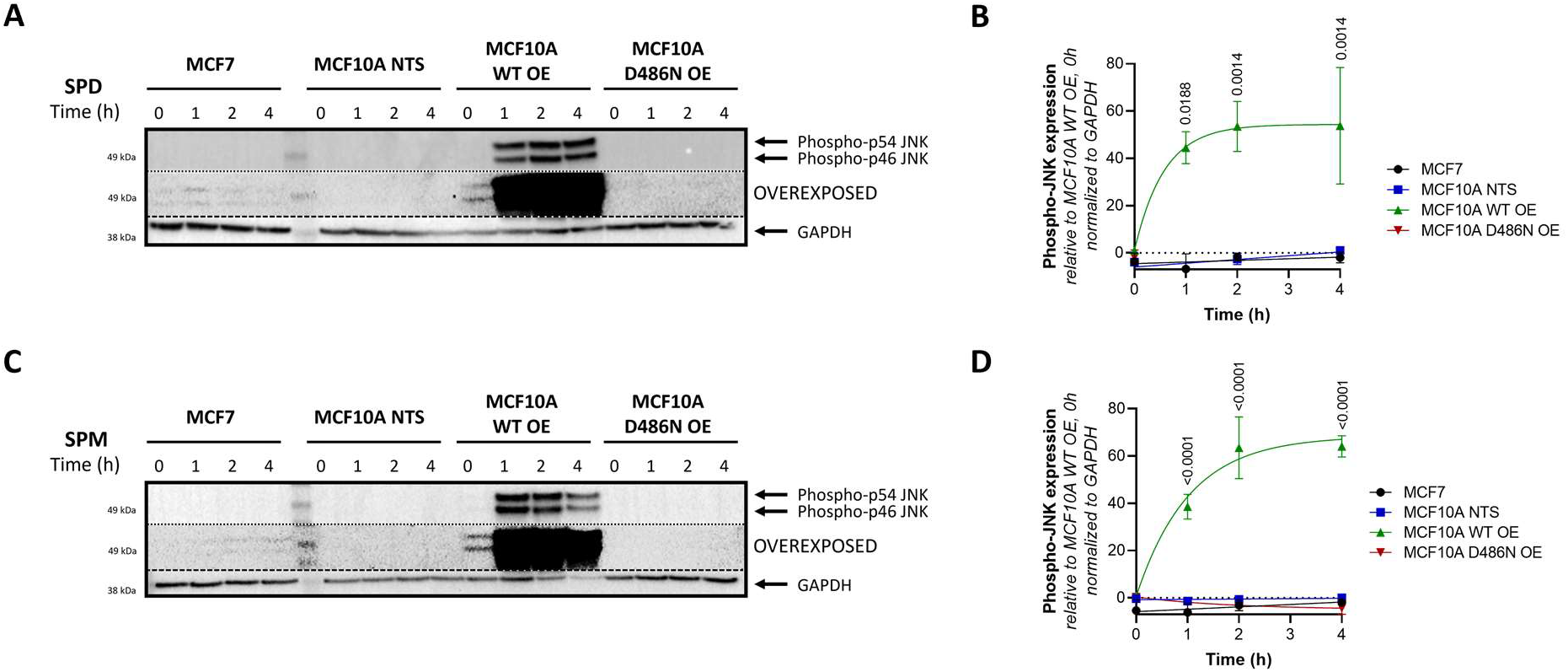
Overexpression of ATP13A4 WT induces JNK phosphorylation in MCF10A cells. MCF7, non-transduced (NTS) MCF10A cells and MCF10A cells overexpressing ATP13A4 WT (WT OE) or a catalytically dead mutant (D486N OE) were treated for indicated times with 100 μM spermidine (**A-B**) or 100 μM spermine (**C-D**). Expression levels of phospho-JNK were determined by Western blot analysis. **A** and **C** depict representative blots of at least three independent experiments. Of note, the blots were overexposed to show the presence of phospho-JNK bands in MCF7 cells. **B** and **D** show the bar graphs of the quantification of phospho-JNK levels. Data were normalized to MCF10A WT OE, untreated control and are presented as the mean ± s.e.m.. Statistical significance was determined by two-way ANOVA with Dunnett’s multiple comparisons test, comparing the different time points to untreated control for each cell line. *p* values are depicted in the graphs, non-significant values are not indicated.

Together, our findings suggest that the ATP13A4-mediated polyamine response is able to modulate the JNK signaling pathway in a time dependent manner.

## 4. Discussion

In this study, we present evidence that the elevated PTS in the commonly studied breast cancer cell line MCF7 may depend on an increased expression of ATP13A4, which we here established as a novel polyamine transporter in the PTS that drives polyamine-dependent phenotypes in breast cancer cells.

### 4.1. ATP13A4 emerges as a novel polyamine transporter in the mammalian PTS

MCF7 cells display upregulated polyamine transport activity in comparison to MCF10A cells, which most likely is attributed to the more than 10-fold higher expression levels of ATP13A4, a previously unexplored member of the P5B-ATPases. Furthermore, we found that overexpression of catalytically active ATP13A4 results in an increased cellular polyamine uptake in MCF10A cells. These data provide the first evidence that ATP13A4 functions as a polyamine transporter in the mammalian PTS, similar to ATP13A2 and ATP13A3. The overlap in polyamine transport function among P5B-ATPases fits well with their close evolutionary relationship, conserved biochemical behavior of spontaneous autophosphorylation, their overlapping endo-/lysosomal localization, peculiar N-terminal topology and predicted three-dimensional structures (AlphaFold) [1, 26]. In addition, all P5B-ATPases present a highly conserved substrate binding site that has been resolved in human ATP13A2 [27-30] and yeast Ypk9p [31] cryo-EM structures as a polyamine binding pocket.

Fluorescently labeled polyamines are genuine substrates of P5B-ATPases stimulating their ATPase activity [19]. They behave remarkably similar as radiolabeled polyamines for ATP13A2 or ATP13A3-dependent uptake in cells, although subtle differences have also been observed [19]. This suggests that the relative uptake capacity towards BODIPY-labeled polyamine analogs is informative, but not conclusive for deducing the substrate specificity of ATP13A4. Our data in the MCF7 and MCF10A cell models with ATP13A4 expression suggest that ATP13A4 presents a broad polyamine specificity, similar to what has been described for ATP13A3 [3, 19]. Indeed, we observed ATP13A4- transport dependent BODIPY-putrescine, -spermidine and -spermine uptake that is competitive with MGBG uptake, a toxic polyamine analog. The close similarity of ATP13A4 and ATP13A3 is in line with their evolutionary relationship. ATP13A4 emerged in higher vertebrates following an ATP13A3 gene duplication in the evolution of lobe-finned fish [26]. However, to unequivocally establish the polyamine-specificity of ATP13A4, an in-depth biochemical characterization using purified enzyme should be carried out. Whereas our results suggest that BV inhibits the PTS activity in MCF7 cells, BV did not abolish polyamine uptake in MCF10A cells overexpressing WT ATP13A4. This points to ATP13A4-independent effects of BV that may be cell type specific, possibly involving upstream BV- sensitive components within the endocytic pathway.

In contrast to the ubiquitously expressed ATP13A2-3 isoforms, ATP13A4 expression is tissue-specific and found mainly in epithelial glandular tissue such as the mammary glands [26], which fits well with a putative role in breast cancer. ATP13A4 has been described in early, recycling and late endosomes, overlapping to some extent with the ATP13A2 and ATP13A3 localization [26]. Polyamines are first endocytosed via heparan sulphate proteoglycans [32] and are subsequently transported via various endosomal P5B ATPase isoforms towards the cytosol. Our results indicate that ATP13A4 may be (partially) redundant in function to ATP13A2 and ATP13A3, but may play a specific role in glandular cells such as mammary gland to further boost PTS activity.

### 4.2. Is ATP13A4 implicated in the upregulated PTS of other cancer types?

Besides a postulated physiological function in glandular tissues, ATP13A4 emerges as a polyamine transporter that is upregulated in breast cancer cells where it contributes to an increased PTS activity. Many cancer types rely on an increased PTS, but it is unlikely that ATP13A4 may be the only P5B- ATPase that is implicated in cancer. Indeed, ATP13A2 has been implicated in several human cancers, including melanoma [33], colon cancer [34], hepatocellular carcinoma [35], acute myeloid leukemia [36] and non-small cell lung cancer [37]; while ATP13A3 has been linked to pancreatic cancer and colorectal cancer [38-40]. Interestingly, high expression of ATP13A3 was found in metastatic pancreatic cancer cells with high polyamine uptake compared to slowly proliferating cells with low import activity [40]. Therefore, dependent on the cancer type, one or more P5B-type ATPase isoform may be upregulated to elevate the PTS in cancerous cells.

So far, two other studies have highlighted a role for ATP13A4 in human cancer. The first study found an association of ATP13A4 with high-grade serous ovarian carcinoma [41], while a second study linked ATP13A4 to lung adenocarcinoma through anaplastic lymphoma kinase (ALK) rearrangements [42]. To further explore the putative role of ATP13A4 in other cancer types, we investigated the prevalence of ATP13A4 genetic alterations across various human cancers using the online database cBioPortal [43] in a pan-cancer dataset [44]. Patients with non-small cell lung cancer showed the highest frequency of ATP13A4 alteration (> 60 %) with amplification as the primary genetic alteration type (Figure 5A). Of note, ATP13A4 underwent amplification in over 10 % of patients with ovarian cancer, cervical cancer, head and neck cancer, endometrial cancer, uterine endometrioid carcinoma, bladder cancer, melanoma, lung cancer and breast cancer. In addition, a potential correlation between ATP13A4 genetic alterations and the clinical survival prognosis of patients was detected. Across cancers, the median overall survival time was significantly shorter for patients with ATP13A4 alterations (Figure 5B, 29.1 months *versus* 57.9 months for reference group). Altogether, these findings confirm that ATP13A4 alterations may be involved in cancer. Further studies will be required to firmly establish the role of ATP13A4 in these other cancer types and to validate ATP13A4 as a candidate therapeutic target to block the PTS in specific cancer types.

**Figure 5.**
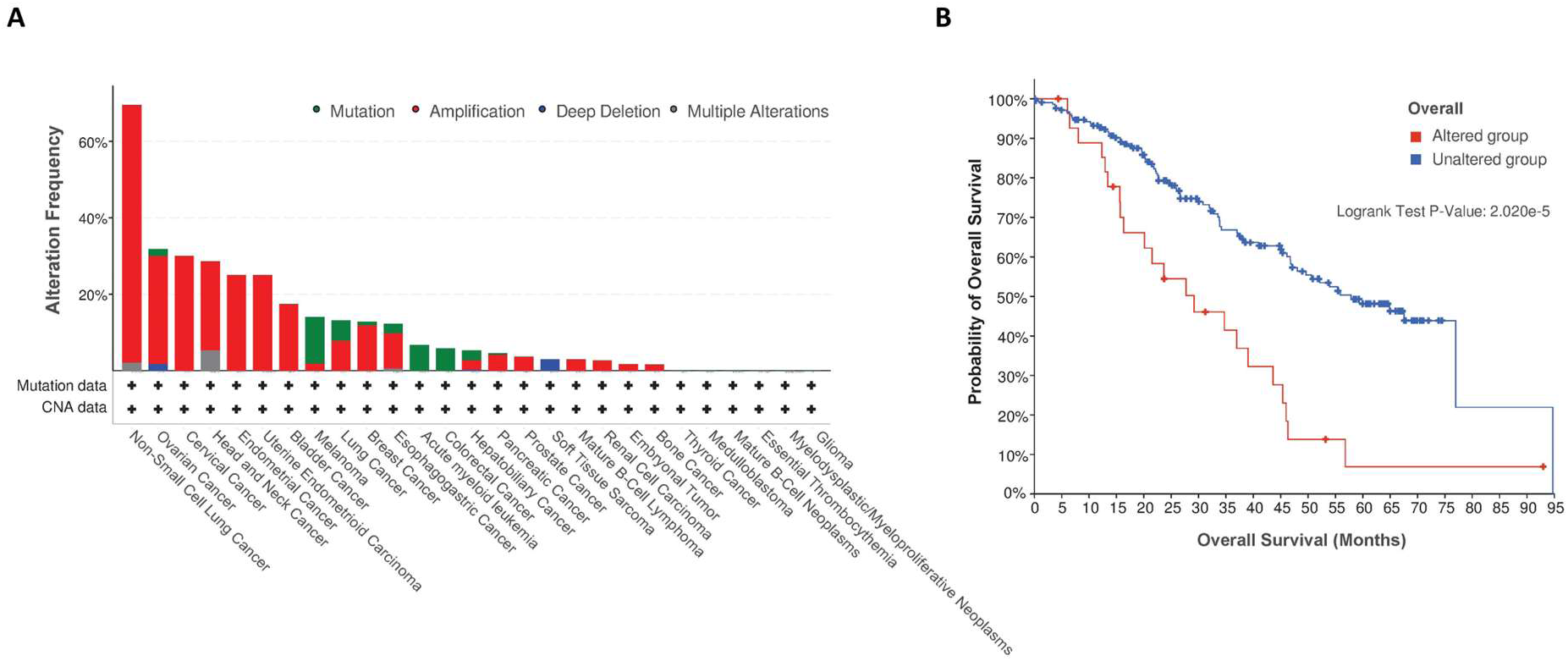
ATP13A4 expression in cancer. **A**. Alteration frequency of ATP13A4 across different cancers from a pan-cancer dataset [44] in cBioPortal [43]. **B**. Overall survival analysis in cases with or without ATP13A4 alterations from the same pan-cancer dataset. Survival analysis significance was based on the log-rank test.

### 4.3. Interplay of ATP13A4, polyamine toxicity and JNK signaling

The ATP13A4-mediated upregulation of the PTS sensitizes cells to polyamine cytotoxicity, which is a surprising finding, since cancer cell proliferation relies on elevated polyamine levels. However, our results are in line with a previous study that reported spermidine-induced apoptosis in cervical cancer [45]. We hypothesize that at high exogenous polyamine concentrations the overactive PTS may lead to harmful intracellular polyamine accumulation that disrupts signaling, polyamine homeostasis and/or transcription and translation [46, 47].

Polyamine accumulation may lead to excessive polyamine degradation with the formation of harmful reactive aldehydes and H2O2 [48]. However, the inhibition of the main degradative enzyme SAT1 with berenil had no differential impact on MCF7 *versus* MCF10A cells, indicating that the polyamine toxicity may occur independently from SAT1. Instead, the observed polyamine cytotoxicity via ATP13A4 may be mediated by activation of the JNK pathway. We found that spermidine and spermine triggered rapid upregulation of JNK phosphorylation in MCF10A cells that overexpress catalytically active ATP13A4, whereas MCF7 cells exhibit high basal levels of JNK phosphorylation that respond to a lesser extent to polyamine supplementation. This difference in magnitude may be explained by the high overexpression of ATP13A4 following lentiviral transduction of the MCF10A cells compared to the endogenous ATP13A4 levels in the MCF7 cells. Of interest, the JNK pathway has been implicated in various cancers and was demonstrated to have both pro-tumorigenic and tumor-suppressive roles in breast cancer [49]. On the one hand, JNK signaling prevents tumor initiation and development in breast cancer [50, 51], whereas on the other hand JNK activity promotes breast cancer metastasis [52, 53] and contributes to tumor aggressiveness via forming an immunosuppressive tumor microenvironment [54]. It was demonstrated that JNK regulates both pro-apoptotic as pro-survival pathways, with prolonged activation of JNK being associated with apoptosis and transient JNK activation with cell proliferation or survival [49]. ATP13A4-mediated polyamine transport may lead to sustained JNK activation with downstream apoptotic consequences.

Our observations suggest that supplementation of polyamines to specifically induce cytotoxicity in cancer cells may be considered as a therapeutic approach, but this may not be without risk. Spermine/spermidine supplementation via the drinking water appears beneficial for longevity of model organisms without inducing cancer, but whether sufficiently high polyamine levels can be reached to induce toxicity in cells with elevated PTS remains questionable. Furthermore, direct injection of polyamines in rodents is lethal [55], indicating that boosting plasma polyamine levels to target cancer cells may cause adverse effects. A possible safer alternative may be the use of toxic polyamine conjugates that preferentially enter cancer cells due to their upregulated PTS, which may be effective in inducing cell death at low concentrations [8].

In conclusion, our findings shed light on ATP13A4 as a novel member of the enigmatic PTS that is implicated in the upregulated PTS in breast cancer cells. Future in-depth characterization of both the cytosolic and organellar impact of ATP13A4’s transport function will be instrumental in further understanding its role in (breast) cancer and may validate ATP13A4 as a candidate therapeutic target for anticancer drugs that block the PTS.

## Supplementary Material

**Figure S1. mRNA levels of P5B ATPase isoforms in MCF7 and MCF10A cell lines**. Boxplots obtained from ARCHS4 [23], a web resource of processed RNA-seq data, showing mRNA expression levels of **(A)** ATP13A4 in different cell lines and **(B)** P5B ATPases ATP13A2-5 in MCF7 *versus* MCF10A cells.

**Figure S2. Benzyl viologen reduces polyamine uptake activity in MCF7 cells**. The impact of benzyl viologen (BV) on cellular uptake of BODIPY-labeled putrescine (BODIPY-PUT) **(A)**, spermidine (BODIPY-SPD) **(B)** and spermine (BODIPY-SPM) **(C)** (1 μM, 2h) in MCF7 *versus* MCF10A cells was assessed via flow cytometry. MFI, mean fluorescence intensity. **(D)** MCF7 and MCF10A cells were treated for 48 h with indicated concentrations of benzyl viologen (BV). Cell viability was assessed via MUH assay and data were normalized to untreated control for each cell line. Data are presented as the mean of a minimum of three independent experiments, with individual data points shown **(A-C)** or ± s.e.m. **(D)**. Statistical significance was determined by one-way ANOVA with Šídák’s multiple comparisons test **(A-C)** or by two-way ANOVA with Šídák’s multiple comparisons test **(D)**. *p* values are depicted in the graphs. Non-significant values are not indicated in **D**.

**Figure S3. Cytotoxicity of polyamine metabolism inhibitors in MCF7 and MCF10A cells. (A)** Overview of polyamine metabolism with pharmacological inhibitors indicated in red. MCF7 and MCF10A cells were treated for 1 week with indicated concentrations of DFMO **(B)**, 4MCHA **(C)**, APCHA **(D)**, MGBG **(E)** and berenil **(F)**. Cell viability was assessed via MUH assay. Data were normalized to untreated control for each cell line and are presented as the mean ± s.e.m. of a minimum of three independent experiments. Statistical significance was determined by two-way ANOVA with Šídák’s multiple comparisons test. *p* values are depicted in the graphs, non-significant values are not indicated. ORN: ornithine; PUT: putrescine; SPD: spermidine; SPM: spermine; ODC: ornithine decarboxylase; SRM: SPD synthase; SMS: SPM synthase; AdoMetDC: S-adenosyl-L-methionine-decarboxylase; SAT1: SPD/SPM- N(1)-acetyltransferase; DFMO: D,L-alpha-difluoromethylornithine; 4MCHA: trans-4- methylcyclohexylamine; APCHA: N-(3-aminopropyl)cyclohexylamine; MGBG: methylglyoxal bis- (guanylhydrazone).

**Figure S4. Benzyl viologen does not impact MCF10A cells with ATP13A4 overexpression**. The impact of benzyl viologen (BV) on cellular uptake of BODIPY-labeled putrescine (BODIPY-PUT) **(A)**, spermidine (BODIPY-SPD) **(B)** and spermine (BODIPY-SPM) **(C)** (1 μM, 2h) in MCF10A cells (non-transduced, NTS; ATP13A4 WT overexpression, WT OE; ATP13A4 D486N mutant overexpression, D486N OE) was assessed via flow cytometry. MFI, mean fluorescence intensity. **(D)** MCF7 and MCF10A cells were treated for 48 h with indicated concentrations of benzyl viologen (BV). Cell viability was assessed via MUH assay and data were normalized to untreated control for each cell line. Data are presented as the mean of a minimum of three independent experiments, with individual data points shown **(A-C)** or ± s.e.m. **(D)**. Statistical significance was determined by one-way ANOVA with Šídák’s multiple comparisons test **(A-C)** or by two-way ANOVA with Šídák’s multiple comparisons test **(D)**. *p* values are depicted in the graphs. Non-significant values are not indicated in **D**.

**Figure S5. Effects of polyamines on phosphorylation of ERK1/2, JNK and AKT in MCF7 *vs*. MCF10A cells**. MCF7 and MCF10A cells were treated for indicated times with 100 μM spermidine **(A-D)** or 100 μM spermine **(E-H)**. Expression levels of phospho-JNK, phospho-ERK1/2 and phospho-AKT were determined by Western blot analysis. **A** and **E** depict representative blots of at least two independent experiments, **B-D** and **F-H** show the bar graphs with the quantification. Data were normalized to MCF7, untreated control and are presented as the mean ± s.e.m.. Statistical significance was determined by two-way ANOVA with Dunnett’s multiple comparisons test, comparing the different time points to untreated control for each cell line. Non-significant values are not indicated.

## Author Contributions

Conceptualization, S.v.V. and P.V.; methodology, S.v.V., J.V.A. and C.V.d.H.; formal analysis, S.v.V.; resources, C.V.d.H.; investigation, S.v.V., A.K., E.A., J.V.A. and C.V.d.H.; writing— original draft preparation, S.v.V.; writing—review and editing, S.v.V. and P.V.; visualization, S.v.V.; supervision, S.v.V. and P.V.; funding acquisition, P.V. All authors have read and agreed to the published version of the manuscript.

## Institutional Review Board Statement

Not applicable.

## Informed Consent Statement

Not applicable.

## Data Availability Statement

The data present in the current study are available from the corresponding authors on reasonable request.

## Acknowledgments

We would like to thank Marina Crabbe and Nathalie Jacobs for technical support; Dr. S. Verhelst (KU Leuven) for the production of fluorescent BODIPY polyamines; and Dr. P. Van Veldhoven (KU Leuven) for the kind gift of MGBG. We also acknowledge our frequent use of the facilities and equipment of the Cell and Tissue Imaging Cluster (Dr. P. Vanden Berghe, KU Leuven), and the FACS Core (Dr. S. Schlenner, KU Leuven). S.v.V. thanks S. Martin for his continuous support and insightful discussions.

