## Supplementary figures and images for "ATP13A4 upregulation drives the elevated polyamine transport system in the breast cancer cell line MCF7"

### Figure S1

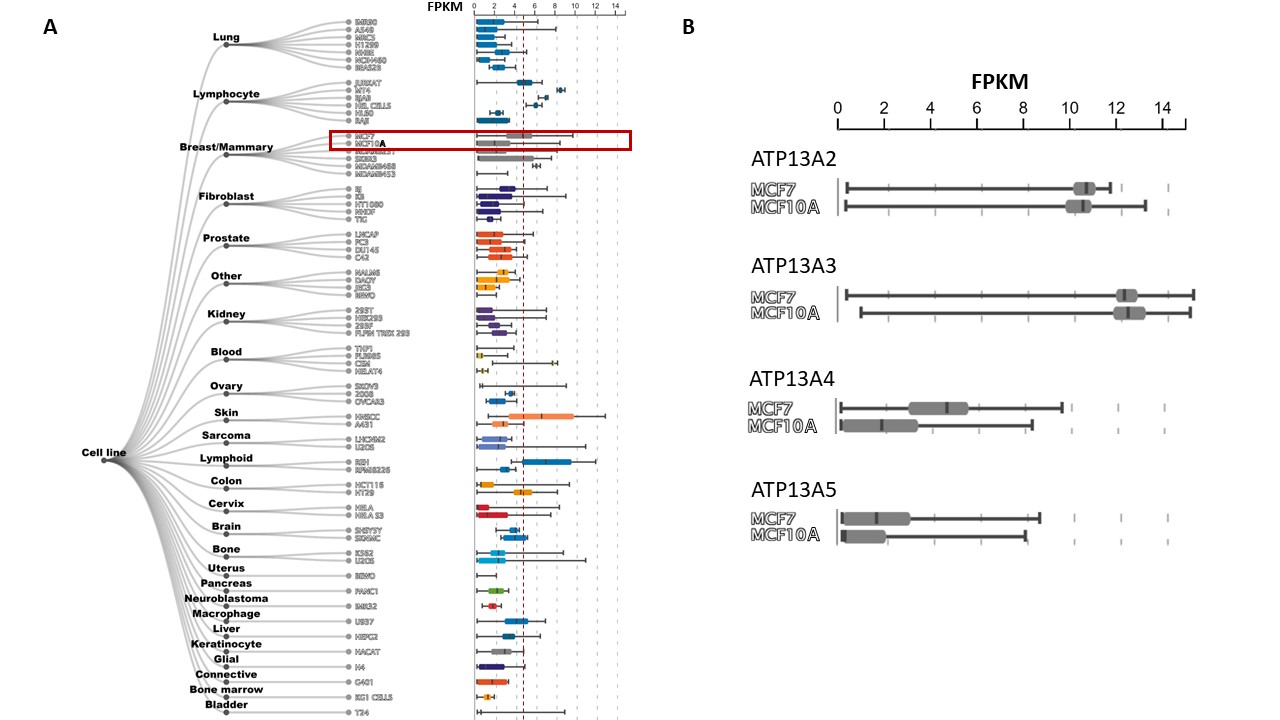

### Figure S2

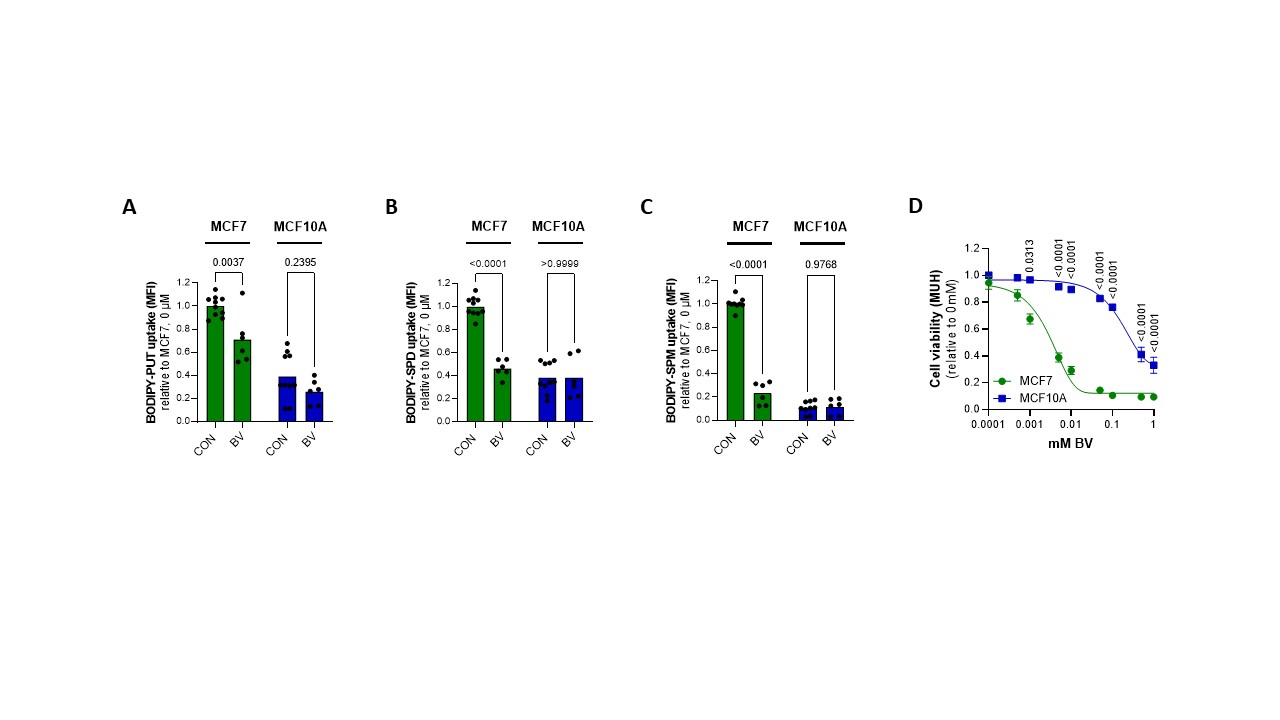

### Figure S3

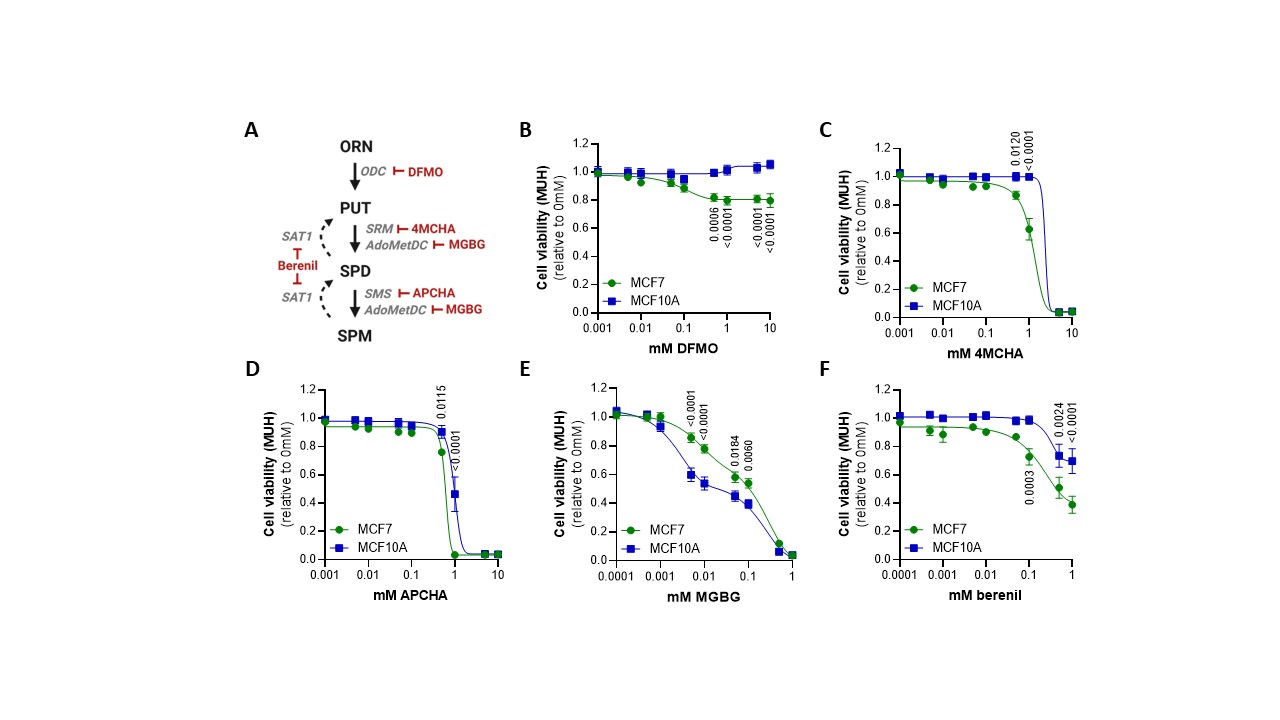

### Figure S4

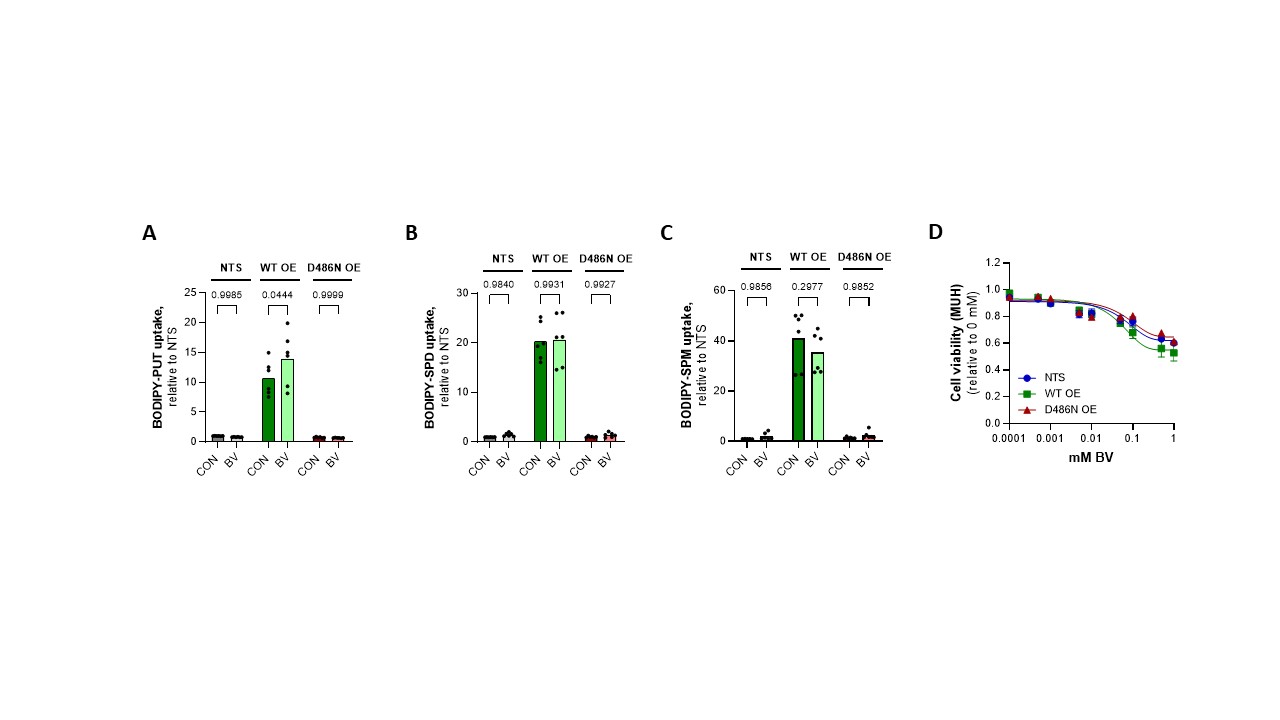

### Figure S5

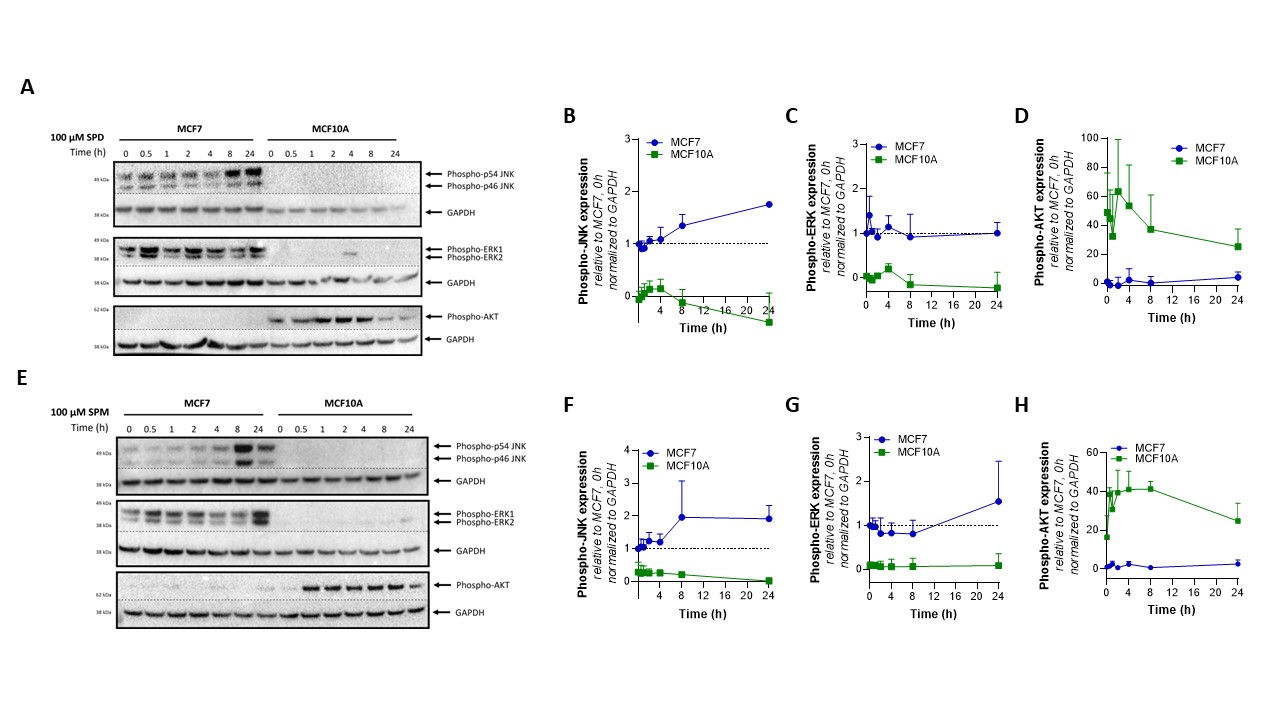
